# The Maximum Entropy Principle For Compositional Data

**DOI:** 10.1101/2022.06.07.495074

**Authors:** Corey Weistuch, Jiening Zhu, Joseph O. Deasy, Allen R. Tannenbaum

## Abstract

In this work, we provide a general method for inferring the stochastic behavior of compositional systems. Our approach is guided by the principle of maximum entropy, a data-driven modeling technique. In particular, we show that our method can accurately capture stochastic, inter-species relationships with minimal model parameters. We provide two proofs of principle. First, we measure the relative abundances of different bacteria and infer how they interact. Second, we show that our method outperforms a common alternative for the extraction of gene-gene interactions in triple-negative breast cancer.

**Author summary:** Compositional systems, represented as proportions of some whole, are ubiquitous. They encompass the abundances of proteins in a cell, the distribution of organisms in nature, and the stoichiometry of the most basic chemical reactions. Thus, a central goal is to understand how such processes emerge from the behaviors of their components and their pairwise interactions. Such a study, however, is challenging for two key reasons. Firstly, such systems are complex and depend, often stochastically, on their constituent parts. Secondly, the data lie on a simplex which influences their correlations. We provide a general and data-driven modeling tool for compositional systems to resolve both of these issues. We achieve this through the principle of maximum entropy, which requires only minimal assumptions and limited experimental data in contrast to current alternatives. We show that our approach provides novel and biologically-intuitive insights and is promising as a comprehensive quantitative framework for compositional data.

## Introduction

Describing the compositions of physical systems, such as in mixtures of industrial chemical reactions, across bacteria in the microbiome, or relative influences in cancer networks is of significant practical importance. In the present work, these systems are modeled as networks of components (or nodes) and their unknown node-node interactions. However, the challenge of inferring these interactions lies in incorporating the defining feature of such compositions: the total proportion across components must always sum to one (or 100%).

Much recent interest has been devoted to improving the statistical analysis of compositional data [1]. The typical strategies that have been employed broadly fall into two categories. First, many apply traditional statistics (such as correlational analyses) Applied to compositional data, however, such tools are known to generate highly inaccurate results [1–3]. The second approach is to consider the simplicial geometry of compositional data more carefully. Here, a simplex describes the set of points that sum to a fixed quantity. Such simplices (triangles, tetrahedrons, etc.) are common in mathematics but less studied in statistics [4, 5]. Current methods for utilizing this information first modify the data (such as through log ratio transformations), but this can introduce unwarranted biases into downstream analysis [1, 6, 7]. What is needed is a more general and principled approach for describing compositional data.

In contrast to previous approaches, we aim to infer a general model for compositional data from first principles. The natural method for this is the principle of maximum entropy or Max Ent [8–11]. Here, one provides constraints, such as means, variances, and even the geometry of the data itself, and Max Ent provides the model. The advantage of this approach is twofold. First, as opposed to other modeling approaches, Max Ent makes minimal assumptions that are not warranted by the data itself [12]. Second, Max Ent is a widely and successfully utilized modeling framework for complex biological systems [13–18]. We provide theory and practical demonstrations of our new approach in the present work.

## The model

Suppose one is given several stochastic observations of the relative abundances of *N* different components. Each of these observations may be represented as a vector Γ = {*s*_1_, *s*_2_,… *s_N_*}. Our goal is to infer the most likely and least-biased inter-component relationships that give rise to these observations (see Fig 1). The unique model with this property is provided by the *principle of maximum entropy,* which selects the model *P* that both maximizes the entropy *S* = – Σ_Γ_ *P*_Γ_ log *P*_Γ_ and satisfies known constraints from the data. Here, the standard constraints are the estimated first and second moments, *M_i_* = 〈*s_i_*〉 and *χ_ij_* = 〈*s_i_s_j_*〉 [19], as well as the special compositional constraint, Σ_*i*_ *s_i_* = 1 (or 100%). The resulting solution *P*\*, obtained through the method of Lagrange multipliers, is given by:

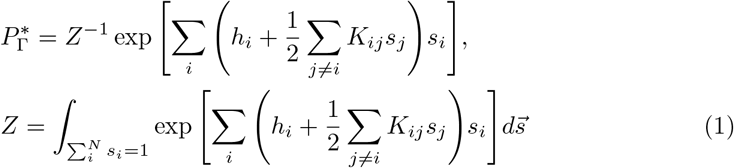

Here *h_i_* and *K_ij_* enforce, respectively, the means *M_i_* and the covariances *η_ij_ – M_i_M_j_*. The normalizing constant *Z* is defined by an intractable integral over the simplex. Thus, the model parameters are found using an adapted pseudolikelihood approximation (see Methods: The simplex pseudolikelihood method). Finally, as Σ_*i*_. *s_i_* = 1, several constraints are redundant. Thus, we set *h_N_* = 0 and *K_ii_* (*i* = 1, 2,… *N*) to 0 (see Methods: Refining the maximum entropy parameters).

**Fig 1.**
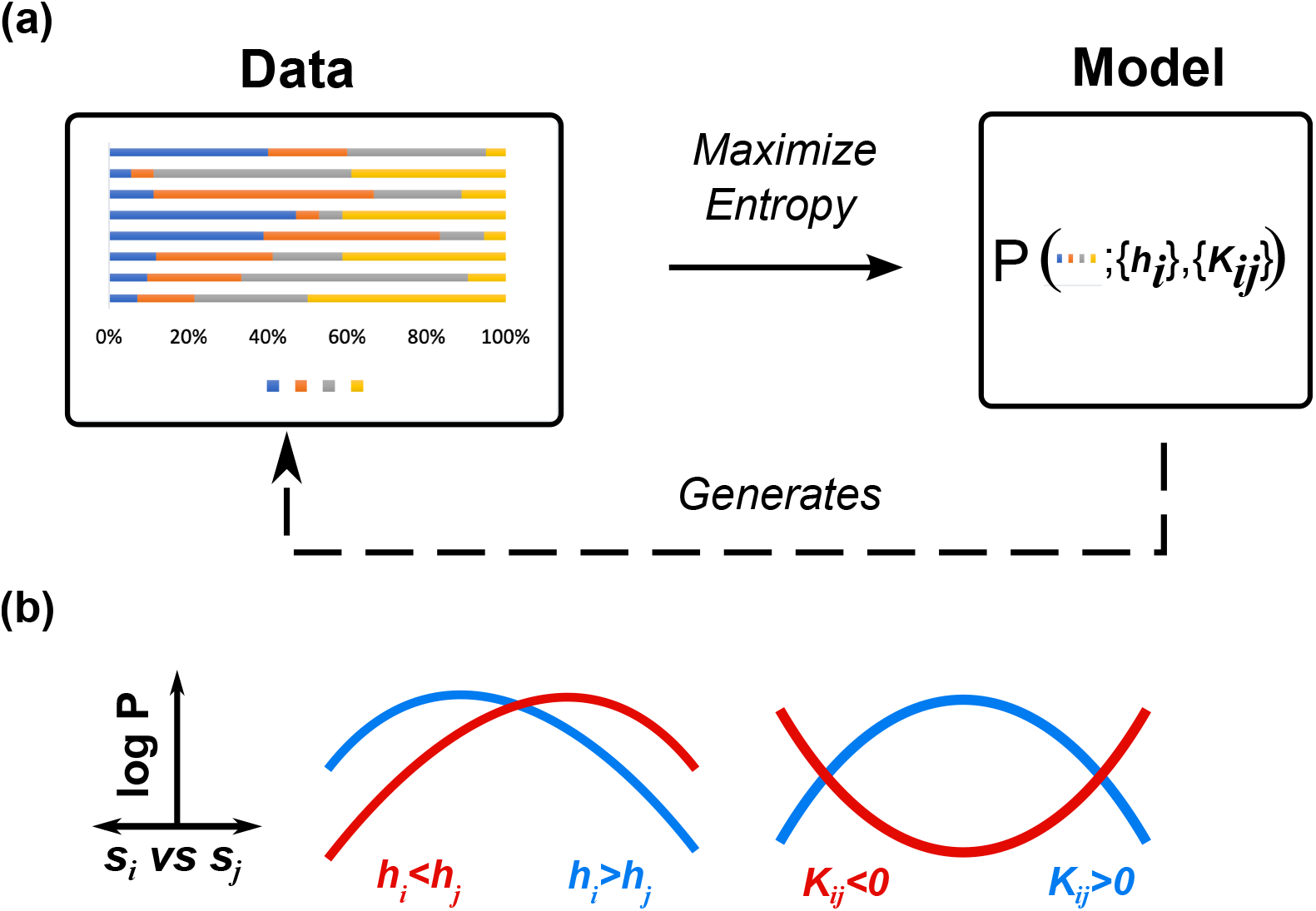
The Compositional Maximum Entropy (CME) approach. (a). Through maximum entropy, CME infers the unknown generative model of the observed component abundances. (b). *h_i_* embodies the influence of each (*i*) component. Components with large *h_i_* tend to have higher abundances than those with small *h_i_*. *K_ij_* embodies the interaction between pairs of components. Pairs with *K_ij_* > 0 tend to coexist, while pairs with *K_ij_* < 0 tend to be mutually exclusive.

In summary, Eq 1 provides the Compositional Maximum Entropy model (CME) subject to known means and covariances. The CME method provides interpretable influence weights *h_i_* for each component node *i* as well as the interaction strengths *K_ij_* between each pair of components (*i* and *j*). Below, we provide two proofs of principle of the method: in a model of the abundances of co-evolving species and the analysis of gene expression data in cancer.

### Quantifying competition among co-evolving species

The quantification of competition among bacteria in the gut, market forces in the economy (or even among scientists) is of course of great interest. A simple and widely-used mechanism is provided by the competitive Lotka-Volterra model (cLV), which describes the population dynamics (i.e., the abundances) of different species vying for a shared resource [20–22]. The population 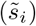 of each species *i* depends on its growth rate *r_i_* and interaction *α_i_* with each other species *j*. Furthermore, the population of each type stops growing as it nears its carrying capacity *κ_i_*, representing the complete exhaustion of resources.

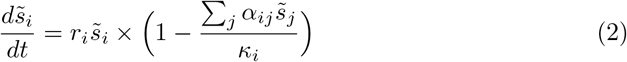

While cLV remains a powerful model for predicting population dynamics, several challenges remain in calibrating it to experimental data. First, we are often only provided with relative (normalized) species abundances. Tools handling both this information loss and the resulting compositional data remain problematic [2, 23]. In addition, we rarely have access to the full time series. Bacterial abundances, for example, are typically measured sparsely but across many different conditions and environments [2].

Here we show that CME can provide accurate quantitative estimates of inter-species interactions, as predicted by cLV, using only available experimental information. The simulated cLV abundances 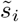 are first normalized to resemble experimental data:

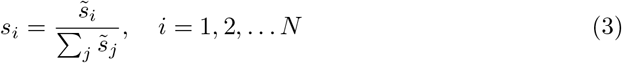

The time-evolving relative abundances *s_i_*(*t*) are then randomly sampled to apply CME. Compared to the cLV model, our proposed approach requires fewer parameters that are thus more resolvable from the limited available data.

cLV models exhibit three broad classes of stable inter-species behaviors: mutualism (they coexist), neutralism (they ignore each other), and competition (only one type can exist at a time) [23]. To illustrate these behaviors, we consider a cLV model of three different species with equal interactions *α_ij_* = *α*. Figure 2 shows the dynamics and abundance distributions for each of three different regimes: *α* = 0.6 (mutualism, Fig. 2a), *α* = 1.2 (neutralism, Fig. 2b), and *α* = 4.0 (competition, Fig. 2c). For simplicity, *r_i_*, *κ_i_*, and the self-interactions *α_ii_* are fixed at 1. Gaussian noise was then added to the simulated dynamics to introduce additional inter-sample variability.

**Fig 2.**
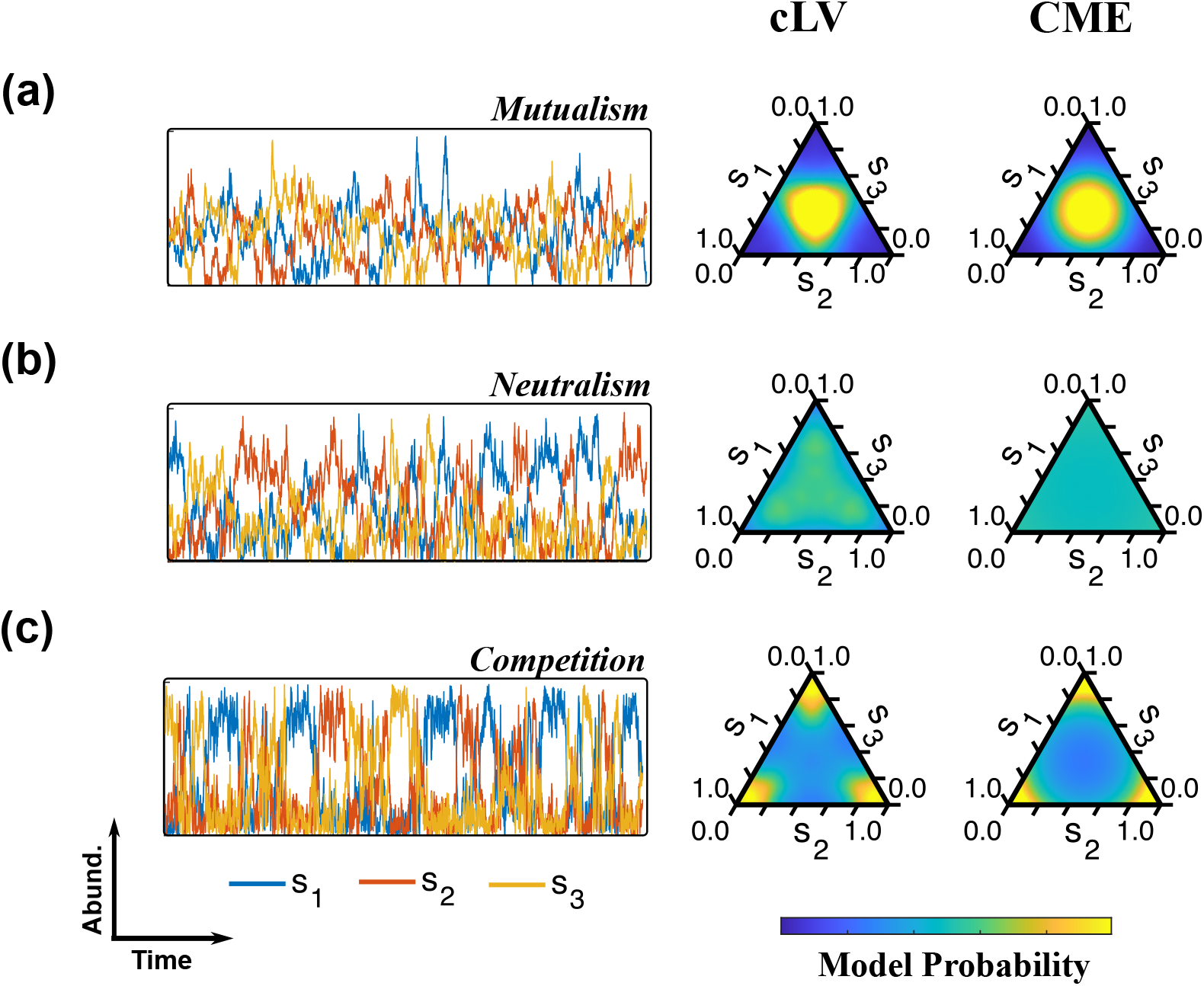
Simulated abundances of three co-evolving species under mutualism (a), neutralism (b), and competition (c). Left, the cLV simulated abundances of each of the three interacting species over time. Center, the corresponding abundance distribution (cLV). Right, the best fit maximum entropy distribution (CME).

The cLV model exhibits a sharp qualitative transformation in its abundance distribution, from a unimodal (Fig. 2a) to a trimodal (Fig. 2c) behavior, as *α* is increased above a critical value (*α* ≈ 1.2, Fig. 2b). Despite requiring fewer parameters (*h* = 0 and *K*, compared to the original four of cLV), CME (right) captures the cLV model behavior (center) across this transformation; *K* > 0 describes mutualism while *K* < 0 describes competition.

In summary, our model provides a simple, data-driven framework for modeling inter-species relationships from limited experimental information. We next consider the more complex case involving heterogeneous interactions from gene expression data in cancer.

### Revealing driving interactions in cancer networks

Cancer is a heterogeneous disease involving complex molecular interactions between many genes. Despite the wealth of information provided by modern experimental tools, the application of such molecular data, including gene expression, to identify effective drug targets continues to face two significant obstacles. First, the accuracy of experimental expression profiles differs between genes [24]. Thus influences from biologically critical but more poorly resolved genes may be overlooked. Second, genes of typical interest often interact, and their effects overlap [25].

Novel network analysis techniques have been developed to refine the genetic signatures of critical genes in cancer. These approaches have been utilized to discover feedback structures in gene interaction networks, identify hubs and bridges, and define measures of robustness and fragility [26–28]. The Wasserstein distance from optimal transport lies as the basis for such methodologies, and in addition to the above references has been directly applied to the stationary (normalized) measures of the networks in question to derive biological information, e.g. showing that pediatric sarcoma data forms a unique cluster [29]. We will now show that CME may provide an important tool for such problems and help point to potential driver genes and their most important interactions.

To test our method, we analyze whole-genome expression data of triple-negative breast tumors, a highly aggressive and complex type of cancer. While many genes are known to be dysregulated in this disease, the relative influence of individual genes is far from established [30]. The data consist of expression profiles from 299 disease samples in METABRIC (Methods: The METABRIC dataset) [31]. We obtained normalized weights for each of *N* = 3147 genes using the Human Protein Reference Database (HPRD) for each sample (Methods: Network identification) [32]. As most of these genes provide no signal in the data, we renormalized these weights after considering only the top 17 highest variabiliy genes with known relevance to cancer (according to OncoKB, see [33] and Methods: Data preparation). Figure 3 illustrates the known connectivity of these genes, but with node size and color proportional to their inferred maximum entropy node weights (*h_i_*). We immediately notice two key details. First, our genes of interest all form a tightly connected network. Second, despite being highly correlated with each other (as the topology would suggest), these genes have unequal influences on the data. The highest-ranked genes, SRC and TP53, are also known master regulators of cancer [34, 35].

**Fig 3.**
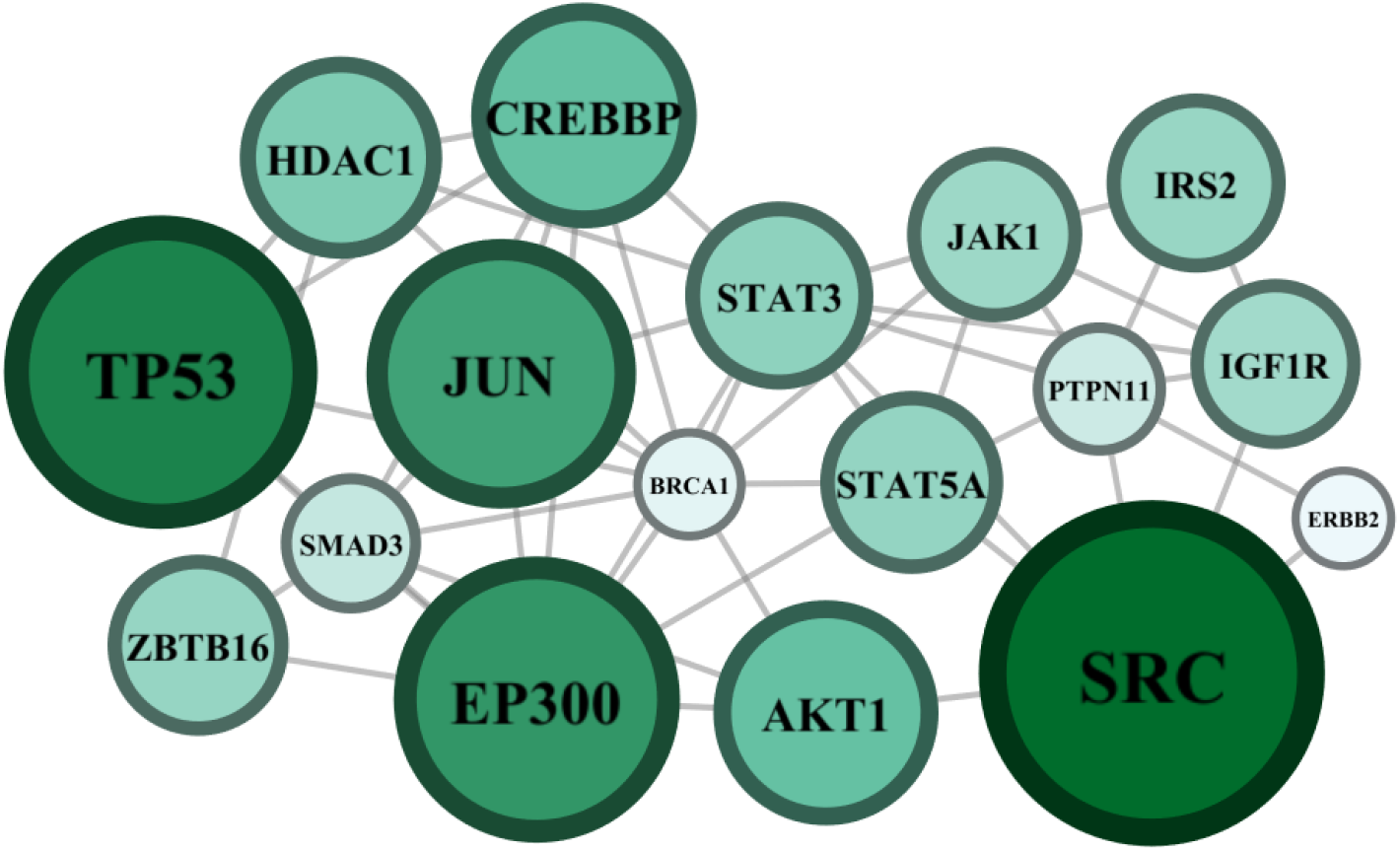
Maximum entropy ranking of key genes in triple-negative breast cancer. Edges correspond to protein-protein interactions obtained from HPRD. Node color and size correspond to their influence (*h_i_*).

A major strength of maximum entropy methods is identifying key node-node interactions underlying the more complex covariances measured from data. This is illustrated in Figure 4, which compares the maximum entropy pairwise interactions *K_ij_* to those inferred from a widely-used alternative statistical model, the logit-normal distribution (Methods: Implementation of the logit-normal distribution); there is an identifiable mapping between the strongest magnitude maximum entropy interactions (Fig. 4a), in contrast to these obtained from the logit-normal (Fig. 4b), and their corresponding gene-gene covariances (Fig. 4c).

**Fig 4.**
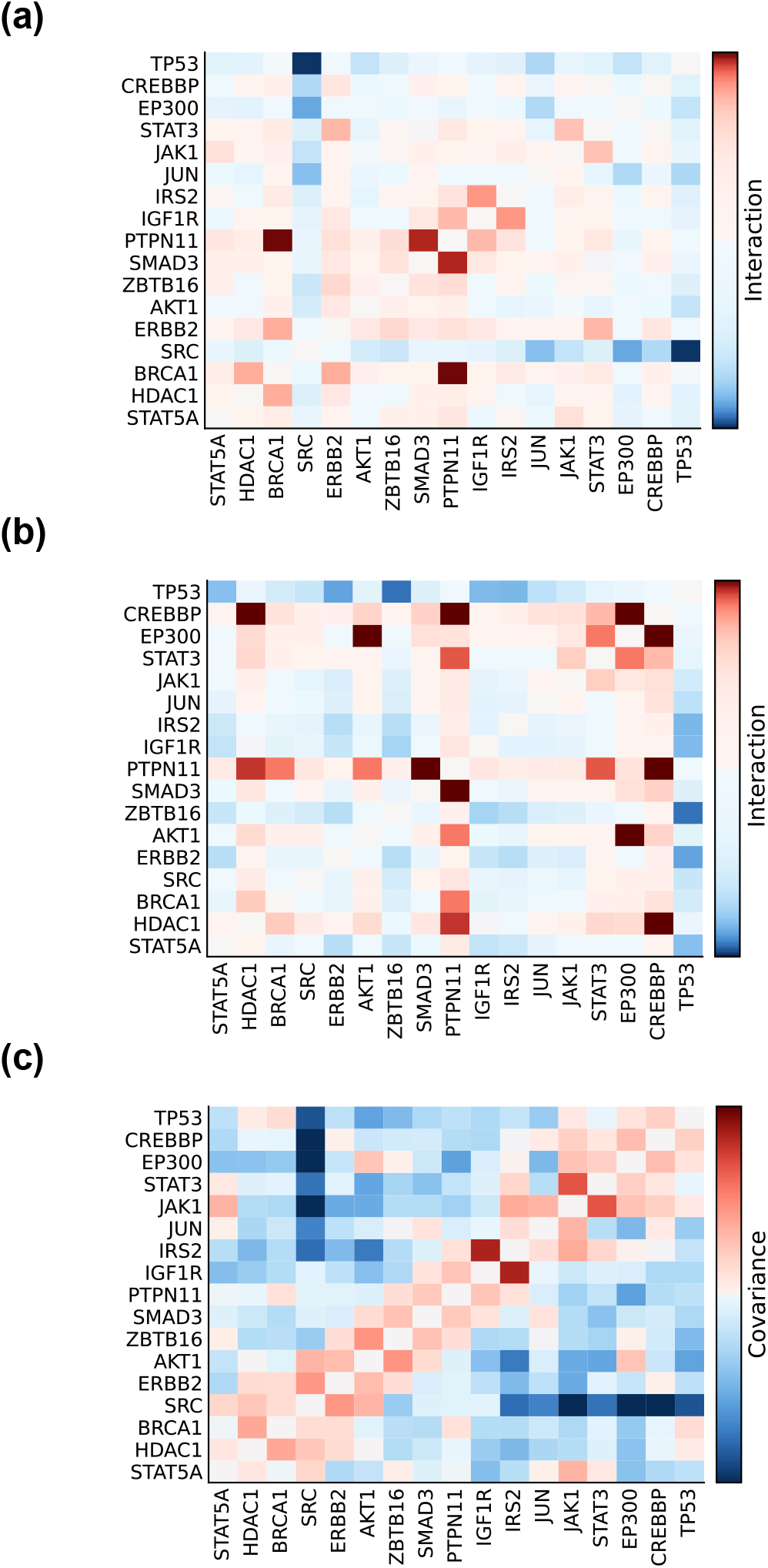
Comparison between three breast cancer network analyses: CME (a), logit-normal (b), and the data covariances (c). Maximum entropy and logit-normal results are shown on a log-scale to reveal the most influential positive (red) and negative (blue) interactions.

We also note that the two top maximum entropy interactions alone (SRC/TP53 and BRCA1/PTPN11) provide an intuitive explanation for some of the key features of the data. SRC and TP53 maintain the critical balance between growth (SRC) and damage repair (TP53): enhanced SRC (or repressed TP53) promotes cell survival, growth, and metastasis, while the reverse leads to accelerated aging [34–36]. This known and critical negative interaction between SRC and TP53 separates most of the 17 genes into two distinct (and negatively covarying) clusters. Thus, since BRCA1 and PTPN11 belong to opposing clusters, their corrected interaction, as revealed by both maximum entropy and logit-normal modeling, is much larger than expected from their weak, positive covariance. Interestingly, both BRCA1 and PTPN11, along with SRC and TP53, are involved in the JAK-STAT pathway [37, 38]. Thus, these genes may have a general and synergistic role in cancer that remains to be explored.

Yet, while the logit-normal model does appear to resolve some features (such as the subtle covariance between AKT1 and EP300) that CME neglects, the interactions predicted by this method generally appear difficult to interpret in the context of the original covariance matrix: it predicts many interactions between uncorrelated genes and fails to resolve, among others, the clear negative covariance between SRC and TP53. Overall, the CME method provides a parsimonious biological mechanism, involving known cancer drivers and only a few of their interactions, for the genetic variability in this poorly understood disease.

## Discussion

We have provided CME, a probabilistic framework for inferring the behaviors of compositional systems from data. Typically, models are deduced bottom-up, starting from mathematical relationships between individual components and combining them often in a complex, nonlinear way. However, as we have described for the Lotka-Volterra model, these interactions can rarely be resolved from the available experimental data. CME, instead, takes a top-down approach – starting from the data and learning the most parsimonious model for it. As evidenced by our breast cancer analysis, CME may also provide more interpretable insights into the organization of compositional systems.

For simplicity, we have considered only small networks; however, our method can be easily extended to much larger networks. First, the pseudolikelihood approach at the core of our method has been successfully applied, with the proper regularization, to networks consisting of thousands of nodes [39]. Second, the implementation of our algorithm uses a scalable L-BFGS algorithm and is fully parallelized across multiple CPU cores.

Similar to partial correlational analysis [40–42], maximum entropy computes direct pairwise interactions by controlling for the confounding indirect effects of the other nodes. Despite being widely used in data analysis and machine learning, partial correlations are only appropriate for linear associations or Gaussian-like data [40]. Maximum entropy methods, such as our application to compositional data [41], are, by contrast, much more general.

Our approach can incorporate more general model constraints as well. The compositional simplex constraint is enforced using the method of Lagrange multipliers. Other geometries [43, 44], even higher-order moments, can be included simply by including new Lagrange multipliers. Finally, as a maximum entropy model, CME is naturally compatible with constraints on other types of data as well [45].

## Conclusion

We proposed CME, a data-driven framework for modeling compositions in multi-species networks. We utilize maximum entropy, a first-principles modeling approach, to learn influential nodes and their network connections using only the available experimental information. Our method requires minimal assumptions and no modifications of the experimental data. Furthermore, the method can be easily generalized to incorporate new types of constraints and data that may emerge.

## Methods

### The simplex pseudolikelihood method

Fitting maximum entropy models to data is generally computationally intractable. Thus, to fit CME, we will adapt the widely-used *pseudolikelihood* approximation [39]. This method requires two pieces of information. First, we need a formula to compute the conditional distribution *P*(*s_i_*|*s*_~*i*_), where *s*_~*i*_ represents all of the variables of interest *S_j_* (*j* = 1, 2,… *N* – 1) excluding *s_i_* and 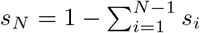. For the simplex model, we have:

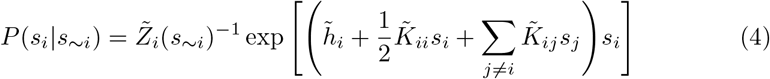

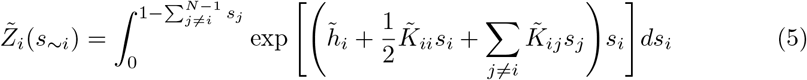

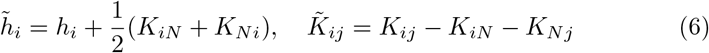

Unlike that of Eq 1, 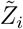 is a tractable Gaussian-like integral. However, its value is sample dependent. Thus, the second required piece of information is the actual samples of the agent proportions 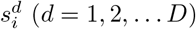 rather than simply the summary means and covariances. Together these enable the maximization of the pseudolikelihood functions 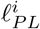 (see Methods: Model implementation):

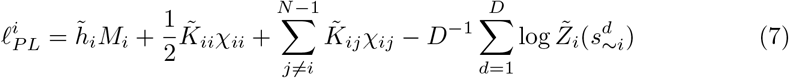

### Refining the maximum entropy parameters

One challenge in modeling compositional data is handling the parameter redundancies induced by the compositional constraint Σ_*i*_ *s_i_* = 1. Specifically, *M_N_*, *χ_iN_*, and *χ_Ni_* are entirely determined from the other data constraints. We could set the associated Lagrange multipliers to 0, but this would hide information about node N (as all of its connections would be forced to 0).

Instead, we recover interpretable model parameters with the following transformations:

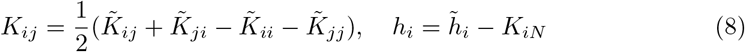

By forcing *K_ii_* to be 0 in Eq 1, we can resolve the interaction strengths between all pairs of nodes in the data. For simplicity, we have defined *h_N_* = 0. However, we can increase or decrease all *h_i_* by any constant and still have an equally good fit. Thus we introduce another transformation to facilitate intra-model comparison of these node weights:

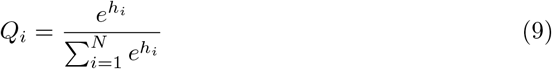

**S1 Fig.**
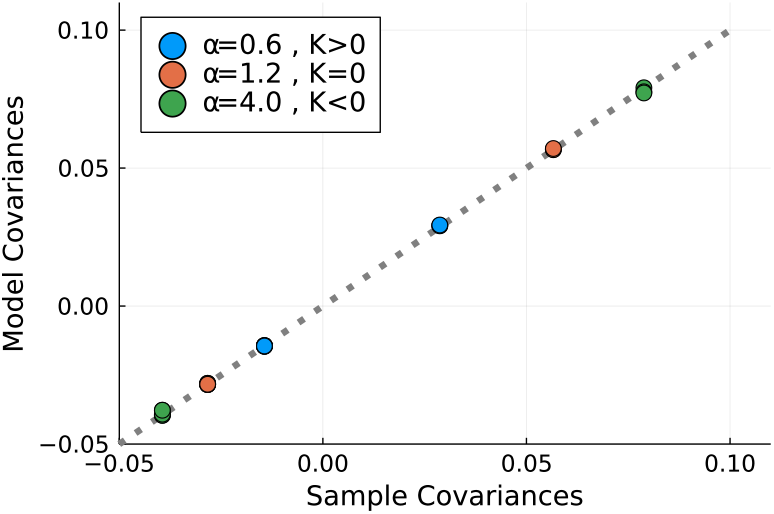
Comparison of CME model covariances to the sample covariances of the cLV model. We observe complete agreement between our model and the data (see Figure 2), confirming the correctness of our maximum entropy fitting algorithm. The means, not shown, were all equal to 1/3 as expected.

Conceptually, the quotient *Q_i_* compares the relative probability of observing a network configuration with influence dominated (*P*\*(*s_i_* = 1)) by node *i*. We posit this as a useful comparison metric for future studies of compositional systems modeled under different conditions.

### Model implementation

To provide a high-accuracy, low overhead approximate maximum of the CME log pseudolikelihood functions, we performed convex optimization using L-BFGS [46] augmented by automatic differentiation. To validate our method, we also designed a custom Monte-Carlo scheme to simulate from CME models. This scheme considers the fitted *h_i_* and *K_ij_* parameters and numerically estimates the corresponding means *M_i_* and covariances Σ_*ij*_ = *χ*_*ij*_ – *M_i_M_j_*. In contrast to CME, such simulation is prohibitively expensive for even moderately-sized, strongly-interacting networks. However, it enabled us to confirm the high accuracy of our model on our Lotka-Volterra simulations (see S1 Figure).

### The METABRIC dataset

Microarray gene expression data for METABRIC were downloaded from the cBioPortal database [47, 48]. The METABRIC dataset, containing 1904 samples, is one of the most extensive publicly-available breast cancer studies [31]. We utilized microarray gene expression data containing 24368 genes from the 299 triple-negative samples.

### Network identification

To quantify the (normalized) influence of genes relevant to triple-negative breast cancer, we utilized the method of network Markov chains [26–28]. The Human Protein Reference Database (HPRD) provides a curated interaction network of most human proteins [32]. Thus, to perform our analysis, we utilized the largest connected component, consisting of 3147 genes, obtained from the intersection of HPRD with the METABRIC gene list. We then performed network analysis as in [28] using the subset of 288 genes annotated in OncoKB, a curated database of prominent cancer genes [33].

### Data preparation

For each sample, we obtain a measure of the relative influence of each of 288 genes. To identify potential drivers of the variability of these influences across the data, we computed their inter-sample Pearson correlations. We identified two distinct clusters of highly correlated genes: one containing a small number of immune-adjacent genes and the other, a much larger component, containing prominent breast cancer genes such as TP53 and BRCA1. Thus, we utilized only this second component for our analysis.

Our primary goal is to identify genes and their interactions that potentially drive the variability in treatment responses observed in triple-negative breast cancer [30]. Likely genes include only those with large influence and inter-subject variability. Upon computing the variance in the influence of each gene, we found 17 candidates with markedly higher variance than the remaining bulk. We thus renormalized node influence across these 17 prime candidates before performing our maximum entropy analysis.

### Implementation of the logit-normal distribution

An alternative to CME, the logit-normal distribution is given by [1]:

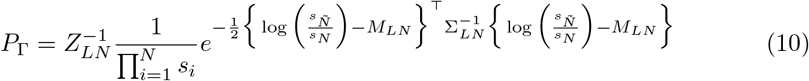

where *M_LN_* and Σ_*LN*_ are the means and covariances of the transformed data: 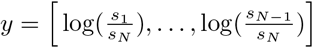. Here, the feature of interest is the precision matrix 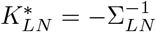 which, under fairly general circumstances, has been shown to approximate maximum entropy interactions [13]. As with CME, we then utilized Eq 8 to define symmetric interactions between all pairs of nodes rather than simply the first *N* – 1.

## Funding

C.W. is funded by the Marie-Josée Kravis Fellowship in Quantitative Biology. A.R.T. is funded by AFOSR grants FA9550-17-1-0435, FA9550-20-1-0029, NIH grant R01AT011419. J.O.D. is funded by NIH grant R21-CA234752 and NIH/NCI Cancer Center Support grant P30 CA008748. J.O.D. and A.R.T. are funded by Breast Cancer Research Foundation Grant BCRF-17-193. The funders had no role in study design, data collection and analysis, decision to publish, or preparation of the manuscript.

## Code availability

The source code and data used to produce the results and analyses presented in this manuscript are available from Github Git repository: https://github.com/Corey651/Compositional_Maximum_Entropy.

## Notes

### Competing Interest Statement

The authors have declared no competing interest.

